# Tendency towards being a “Morning person” increases risk of Parkinson’s disease: evidence from Mendelian randomisation

**DOI:** 10.1101/288241

**Authors:** AJ Noyce, DA Kia, K Heilbron, JEC Jepson, G Hemani, The International Parkinson’s Disease Genomics Consortium, The 23andMe Research Team, DA Hinds, DA Lawlor, Smith G Davey, J Hardy, A Singleton, MA Nalls, NW Wood

**Affiliations:** Preventive Neurology Unit, Wolfson Institute of Preventive Medicine, Queen Mary University of London, London, UK; Department of Molecular Neuroscience, UCL Institute of Neurology, London, UK; 23andMe, Inc., 899 W Evelyn Avenue, Mountain View, California 94041 USA; Department of Clinical and Experimental Epilepsy, UCL Institute of Neurology, London, UK; MRC Integrative Epidemiology Unit at the University of Bristol, Bristol, UK; Population Health Science, Bristol Medical School, University of Bristol, Bristol, UK; Laboratory for Neurogenetics, National Institutes for Health, Bethesda, USA; Data Technica International, USA

**Author notes:** These authors contributed equally (AJ Noyce, DA Kia). **Corresponding author:** Professor Nicholas Wood. Department of Molecular Neuroscience, Institute of Neurology, Queen Square, London WC1N 3BG, UK.

## Abstract

**Background:** Circadian rhythm may play a role in neurodegenerative diseases such as Parkinson’s disease (PD). Chronotype is the behavioural manifestation of circadian rhythm and Mendelian randomisation (MR) involves the use of genetic variants to explore causal effects of exposures on outcomes. This study aimed to explore a causal relationship between chronotype and coffee consumption on risk of PD.

**Methods:** Two-sample MR was undertaken using publicly available GWAS data. Associations between genetic instrumental variables (IV) and “morning person” (one extreme of chronotype) were obtained from the personal genetics company 23andMe, Inc., and UK Biobank, and consisted of the per-allele odds ratio of being a “morning person” for 15 independent variants. The per-allele difference in log-odds of PD for each variant was estimated from a recent meta-analysis. The inverse variance weight method was used to estimate an odds ratio (OR) for the effect of being a “morning person” on PD. Additional MR methods were used to check for bias in the IVW estimate, arising through violation of MR assumptions. The results were compared to analyses employing a genetic instrument of coffee consumption, because coffee consumption has been previously inversely linked to PD.

**Findings:** Being a “morning person” was causally linked with risk of PD (OR 1⋅27; 95% confidence interval 1⋅06-1⋅51; p=0⋅012). Sensitivity analyses did not suggest that invalid instruments were biasing the effect estimate and there was no evidence for a reverse causal relationship between liability for PD and chronotype. There was no robust evidence for a causal effect of high coffee consumption using IV analysis, but the effect was imprecisely estimated (OR 1⋅12; 95% CI 0⋅89-1⋅42; p=0⋅22).

**Interpretation:** We observed causal evidence to support the notion that being a “morning person”, a phenotype driven by the circadian clock, is associated with a higher risk of PD. Further work on the mechanisms is warranted and may lead to novel therapeutic targets.

**Funding:** No specific funding source.

## Introduction

Mendelian randomisation (MR) involves using genetic variants as instruments to study causal relationships between exposures and outcomes.^1^ MR has great potential application in the study of Parkinson’s disease (PD) and other degenerative neurological conditions, given long prodromal phases that may give rise to associations driven by reverse causation in observational studies.^2^ Chronotype is the behavioural manifestation of our internal body clock, which drives circadian changes in a wide range of physiological parameters, including sleep/wake cycles.^3^ Variation in the chronotype underpins whether or not we tend to be “morning persons” or those that function better at night. There is mounting evidence to suggest that circadian rhythm is disrupted in patients with PD and may explain a variety of features of the earliest phases, before diagnosis.^4,5^ Furthermore, in observational studies, individuals that work night shifts appeared to be at lower risk of PD, and those that are intolerant of night time working have been reported to be at higher risk.^6^

Consistent observational study associations have been reported in meta-analyses of coffee consumption and risk of PD, suggesting that those that drink coffee are less likely to be diagnosed with PD.^7^ This association may be causal or may be indicative of a pre-morbid PD personality type, given that negative observational associations have also been described for smoking and alcohol. Using Mendelian randomisation (MR) we sought to determine whether there was evidence that observational associations between morning personality and PD, and/or coffee consumption and PD, were causal.^8,9^

## Methods

Two-sample MR was undertaken using genome wide association (GWA) study data. Two-sample MR involves determining the association between genetic variants and an exposure (for example chronotype) in one sample, and the relationship between those same variants and an outcome of interest (for example PD) in a second sample. From this information, change in the outcome for a given change in the exposure can be estimated, as long as certain assumptions are valid (further details on how MR can be used for causal inference can be found in the supplementary material).^1,10,11^ Ethical approval was not sought for this specific project because all data came from the summary statistics of GWA studies, which were conducted with ethical approval.

### Parkinson’s disease data

The PD GWA study summary statistics used were from a meta-analysis of GWA studies in PD, which related 7,782,514 genetic variants (after imputation) to PD in up to 13,708 PD cases and 95,282 controls from 15 independent GWA datasets of European descent, undertaken by the International Parkinson’s Disease Genomics Consortium (IPDGC;http://www.pdgene.org).^12^

### Morning person (chronotype) instrument

Recently, two GWA studies linking genetic variants to morning or evening chronotype have been reported.^3,13^ The first of these was a GWA study undertaken in 89,283 participants from 23andMe who self-reported whether they considered themselves a “morning person” or not. The phenotype merged answers to two questions with highly correlated answers: 1) ‘Are you naturally a night person or a morning person?’ (Answers: Night owl, Early bird, Neither), and 2) ‘Are you naturally a night person or a morning person?’ (Night person, Morning person, Neither, It depends, I’m not sure). Neutral responses to these two questions were removed.^13^ Fifteen loci reached genome wide significance comparing the extremes of the phenotype (morning person or night person), including seven that were located near established clock genes and an additional four with a plausible role in circadian rhythm (see supplementary material). The second published GWA study of chronotype was undertaken in 100,420 participants from the UK Biobank study, which used the following validated question for assessing chronotype – ‘Do you consider yourself to be… (Answers: ‘Definitely a morning person’, ‘More a morning than evening person, ‘More an evening than a morning person’, ‘Definitely an evening person’, ‘Do not know’, ‘Prefer not to answer’?^3^

IPDGC data contains both cases and controls from 23andMe (see supplementary material). In two-sample MR, where there is overlap between controls in the exposure and outcome dataset, there exists a risk of bias. For this reason, we used the variants (SNPs) in the chronotype loci discovered in the 23andMe data, but instead of using the effect estimates and standard errors from this discovery dataset, we used the replicated effect estimates and standard errors from the UK Biobank data, regardless of the p-value for association. By doing so we mitigated the potential bias due to overlapping samples.

We looked for linkage disequilibrium between variants that were reported as tagging the same chromosome as one another using LD link (https://analysistools.nci.nih.gov/LDlink/). The R^2^ between variants was <0⋅005 in all cases. All 15 chronotype variants were present in the IPDGC data, meaning that there was no need to identify proxies. We extracted the per-allele log-odds ratio of PD together with its standard error (SE) for each of the independent variants. To ascertain the instrument strength, we estimated a first stage F statistic derived from the following formula:

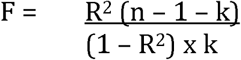

Here, k is the number of variants and n is the number of subjects in the outcome case control study. The combined R^2^ for chronotype variants was estimated using code from MR-Base and the odds ratios, effect alleles frequency and proportion of case/controls from the UK Biobank.^14^ We estimated that the combination of these 15 variants explained 0⋅69% of the variation in chronotype (R^2^ = 0⋅0069).

We repeated the analysis using the effect sizes and standard errors from the 23andMe chronotype GWAS to check whether there were substantial differences between these and the UK Biobank data caused by overlap in the subjects in the exposure and outcome cohort. We also repeated the analysis using unpublished summary statistics from the whole UK Biobank that related to the 15 SNPs in our original instrument (available via the Broad Institute). Finally we generated new instruments from unpublished UK Biobank and 23andMe data to seek further confirmation of the observations made using published instruments.

### Coffee instrument

A GWA study of coffee consumption was published using data from the Coffee and Caffeine Genetics consortium, which was undertaken in 91,462 subjects.^15^ A secondary analysis considered high coffee consumption versus no/low coffee consumption. Six variants in this analysis reached genome wide significance, of which four were in independent loci. Again, all four variants were present in the IPDGC data. We extracted the per-allele log-odds ratio of PD together with its SE for each of these independent variants. We calculated the first stage F statistic for the 4 coffee variants (combined R^2^ = 0⋅015, or 1⋅5% of the variation in coffee consumption).

### Statistical analysis

Two-sample MR was undertaken using previously described methods and as summarised below.^10^ Wald ratios (β_IV_) were calculated for each of the variants in morning person instrument by dividing the per-allele log-odds ratio of PD (β_ZY_) by the log-odds ratio of being a morning person (β_ZX_).

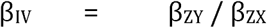

95% confidence intervals (95% CI) were calculated from the SE of each Wald ratio, which was derived from the SE of the SNP-outcome association divided by the SNP-exposure association. Linear regression of the variant-exposure association and variant-outcome association for each instrument was undertaken and weighted by inverse variance. The inverse variance weighted (IVW) method was used to derive a combined point estimate for the odds of PD due to self-identification of being a morning person. The IVW method assumes that all variants are valid IVs (see supplementary material).

Individual Wald ratios and 95% CIs were compiled in a forest plot. Heterogeneity in Wald ratios was tested using Cochran’s Q and quantified using the I^2^ test. MR-Egger was used to demonstrate evidence against an aggregate unbalanced horizontal pleiotropic effect. With MR-Egger, the regression line is not constrained to pass through the origin, and the intercept indicates whether there is a net horizontal pleiotropic effect.^10^ Substantial heterogeneity from the IVW estimate indicates that alternative pathways may exist from some of the SNPs to the outcome (known as horizontal pleiotropy), but this doesn’t necessarily bias the estimate (if it is balanced). However, a non-zero intercept from MR-Egger indicates that there may be net directional bias in the IVW estimates. In performing MR-Egger, the third IV assumption of no horizontal pleiotropy from individual variants can be relaxed, but a new assumption arises. The Instrument Strength Independent of the Direct Effect (InSIDE) assumption of MR-Egger will be violated if the genetic instrument-risk factor association (here the joint association of the 15 SNPs with chronotype) is correlated with any pleiotropic associations from the SNPs to the outcome.^10^

Two further MR methods were used to repeat the analysis, which relax other assumptions of IV analysis. The weighted median (WM) method gives consistent effect estimates under the assumption that no more than 50% of the weight of the MR effect estimate comes from invalid (e.g. pleiotropic) SNPs, where weight is determined by the strength of their association with the risk factor (here chronotype).^16^ The modal estimate was then used as a further sensitivity analysis because it is robust to horizontal pleiotropy, and its power to detect a causal effect is greater than the MR-Egger method.^17^ Its assumption is that the most common causal effect estimate is a consistent measure of the true causal effect, even if the majority of SNPs associated with chronotype are invalid instruments. Given the small number of variants available for coffee consumption, an IVW estimate was calculated, but subsequent sensitivity analyses were not undertaken.

### Reverse causation of chronotype and PD

We used MR to test whether liability towards PD risk was causally related to liability towards being a morning person. Because we had access to individual-level 23andMe data for this analysis, we were able to combine 28 independent PD risk SNPs identified in IPDGC meta-GWAS study into a single PD genetic risk score for each individual, representing a much stronger instrument variable (one sample MR).^12^ Specifically:

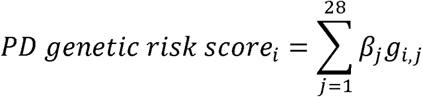

where *i* indicates an individual,*j* indicates one of the 28 PD risk SNPs, β_j_ is the log odds ratio of SNP *j* (taken from the meta-analysis^12^), and *g*_*i,j*_ is the number of effect alleles carried by individual *i* for SNP *j*. Normalised PD genetic risk scores were constructed for 730,872 individuals, none of whom were included in the PD GWA study meta-analysis^12^ (“morning persons”: n=372,909; “evening persons”: n=357,963; people with PD: n=5,201; people without PD: n=725,671). A single Wald ratio and standard error for the PD genetic risk score was calculated using “morning person” as the outcome and PD as the exposure.

### Power calculations

Power calculations were undertaken for the chronotype, coffee consumption, and PD IV analyses.^18^ We had 81% statistical power assuming no heterogeneity between genetic IVs (SNPs) to detect a relative 30% or greater difference in PD risk for a self-identification of being a “morning person” in the IPDGC cases and controls with an alpha of 5% (p-value ≤0⋅05). The statistical power fell to 46% to detect a relative difference of 20% or greater. We had >99% statistical power to detect a 33% difference in PD relative risk for coffee drinkers and 62% statistical power to detect a 16⋅5% difference. For the reverse causation analysis, the R^2^ was 0⋅00079 and the statistic power for a 15% change and 30% change in the outcome respectively, was 39% and 88%.

### Role of the funding source

No specific funding was sought for this work.

## Results

The first stage F statistic for the morning person analysis was 16⋅5. Figure 1 shows a forest plot for the morning person IV analysis, which includes the Wald ratios for individual instruments and the pooled IVW, MR-Egger, weighted median and modal estimates. Using the IVW method, the estimated risk of PD in those identifying as morning persons (compared to those that identified as evening persons) was increased (OR = 1⋅27; 95% CI 1⋅06-1⋅51; p=0⋅012). There was no strong evidence of heterogeneity between Wald ratios for individual instruments (Q=10⋅8, I^2^=0%, p=0⋅705). In the sensitivity analyses, MR-Egger was used to detect net directional pleiotropy but we did not find evidence to suggest that the estimate was biased (intercept −0⋅005; p=0⋅645). The MR-Egger estimate for risk of PD in those identifying as morning persons was OR 1⋅34 (95% CI 0⋅99-1⋅80; p=0⋅058). Further evidence against bias due to invalid instruments came from the weighted median analysis which gave an OR of 1⋅25 (95% CI 1⋅01-1⋅49; p=0⋅066), and from the weighted modal analysis the OR was 1⋅27 (95% CI 1⋅03-1⋅51; p=0⋅075). The point estimates from the three sensitivity analyses all supported a causal association and increased risk of PD in “morning persons”. Using ORs and SEs from the 23andMe chronotype GWAS and unpublished summary statistics from the full-size UK Biobank chronotype GWAS for the same 15 variant instrument yielded qualitatively similar results (see supplementary material). However, new chronotype instruments generated from unpublished UK Biobank GWAS data and 23andMe gave divergent results; the UK Biobank instrument yielded an overall null in the association between chronotype and PD, but may have been biased. A larger 23andMe chronotype instrument supported the results of the main analysis, showing a causal association between chronotype and PD. Both of these additional analyses are discussed in detail in the supplementary material.

**Figure 1.**
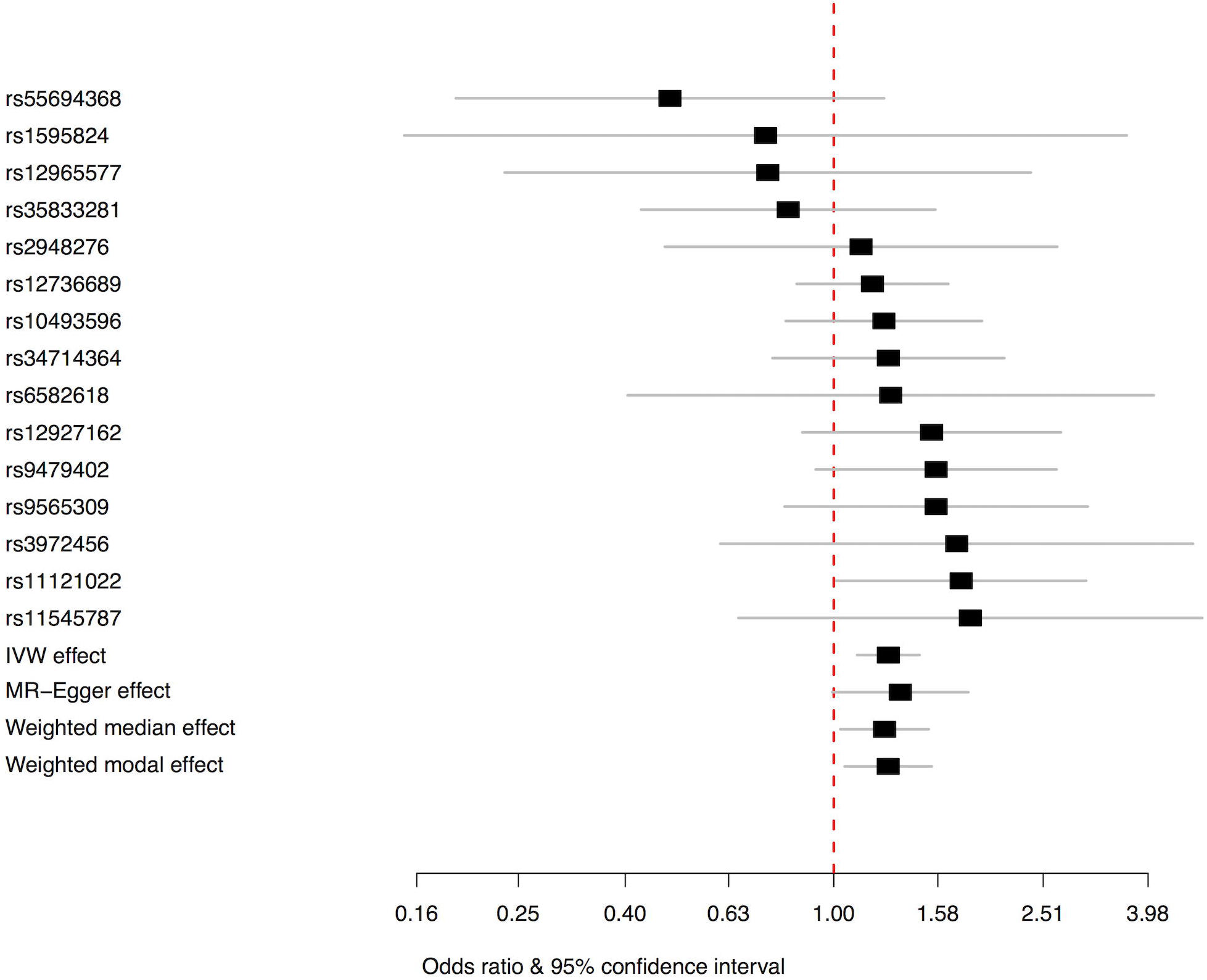
Title: Forest plot of Wald ratios and 95% CIs generated from SNPs associated with chronotype (here being a “morning person”) and risk of PD. Legend: Wald ratios for individual SNPs are listed according to magnitude of effect in the instrumental variable analysis and are presented with pooled effects using the IVW method and sensitivity analyses including MR-Egger regression, and weight median and modal estimate. The squares represent the point estimate and the bars are the 95% confidence intervals.

We tested for reverse causality but this analysis did not suggest that liability towards PD was causal for being a morning person (for a one standard deviation increase in PD genetic risk score: OR 0⋅99, 95% CI 0⋅98-1⋅01, p=0⋅259).

For the coffee analysis, the first stage F-statistic was 179. Figure 2 shows a forest plot for the coffee consumption IV analysis with Wald ratios for each variant and pooled IVW estimates, along with the observational study estimate for the risk of PD associated with coffee consumption (OR 0⋅67; 95% CI 0⋅58-0⋅76).^7^ The estimated risk of PD in those that drank high quantities of coffee relative to those that drank little or none yielded an overall null result in the IVW analysis (OR 1⋅12; 95% CI 0⋅89-1⋅42; p=0⋅22). There was some evidence of heterogeneity in the individual Wald ratio estimates (Q=6⋅9, I^2^=57%, p=0⋅074). MR-Egger was used to detect net directional pleiotropy but there was no convincing evidence to suggest that the estimate was biased (intercept −0⋅052; p=0⋅323). The MR-Egger estimate for risk of PD in those identifying as morning persons was OR 1⋅46 (95% CI 0⋅58-3⋅72; p=0⋅221).

**Figure 2.**
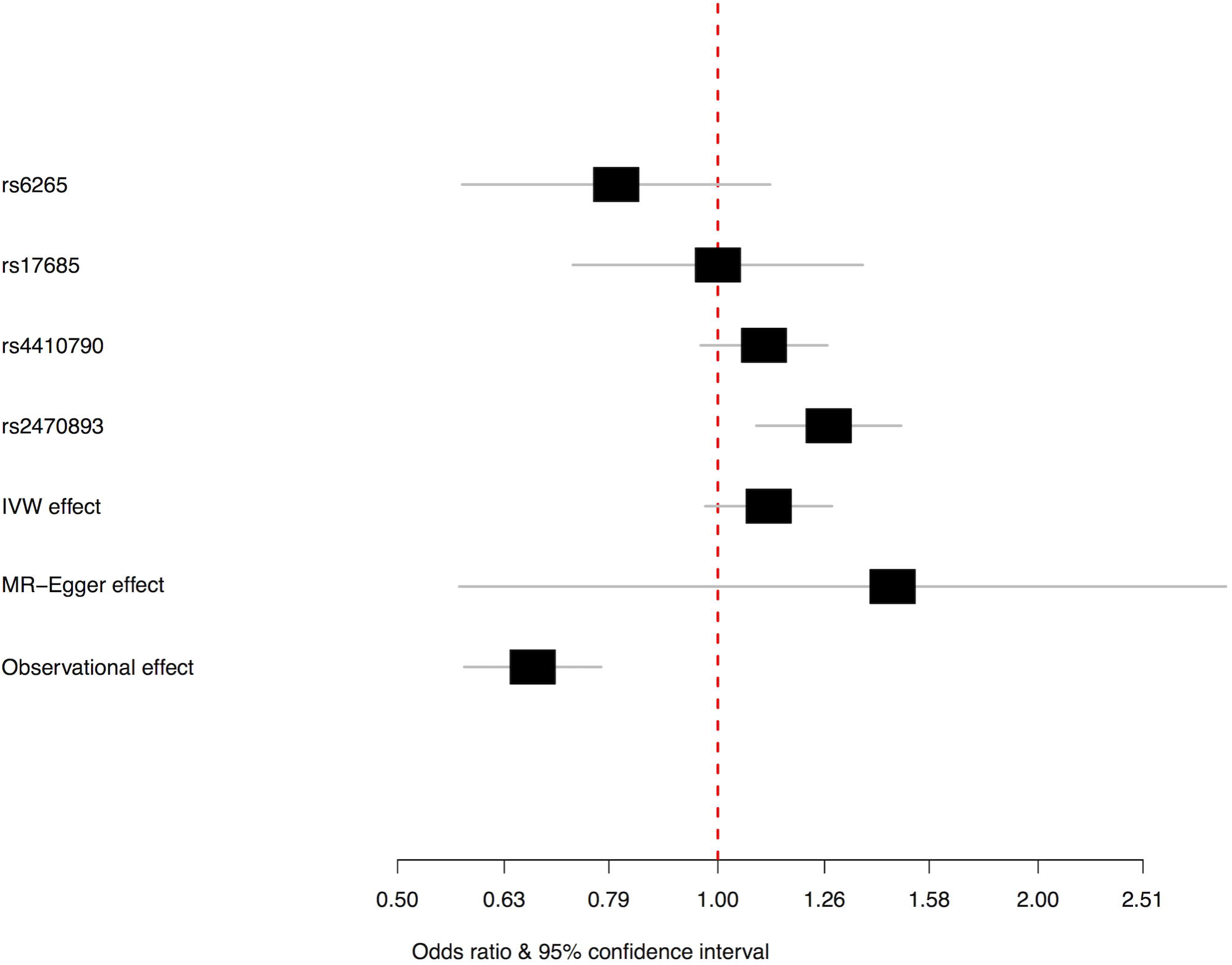
Title: Forest plot of Wald ratios and 95% CIs generated from SNPs associated with coffee consumption (here high compared with low/none) and risk of PD. Legend: Wald ratios for individual SNPs are listed according to magnitude of effect in the instrumental variable analysis and are presented with pooled effects using the IVW method and MR-Egger regression. The observational study association for coffee consumption and risk of PD is also presented and taken from Noyce et al, 2012.^7^ The squares represent the point estimate and the bars are the 95% confidence intervals.

## Discussion

We used two-sample MR to explore a potential causal association between liability towards being a “morning person” and PD. Genetic variants associated with self-reporting as “morning person” (one extreme of chronotype) were used as instrumental variables. We observed evidence of a causal increase in risk of PD in those that identified as being “morning persons”. The magnitude of the effect from the IVW analysis was a 27% increase in risk. Conversely, we found no strong evidence that liability towards increased risk of PD was causal for chronotype (i.e. reverse causation). Our findings lend support to the previous observation that working night shifts was associated with a reduced risk of PD and an increased risk of PD in those that were intolerant of night shifts.^6^

There has been speculation over many years that personality might be linked to PD.^19^ Although this analysis only focuses on one aspect of personality, and one that is regulated by innate circadian mechanisms, it is plausible that other personality traits may also influence risk of PD. Traits such as introspection and conscientiousness were recognised in early observational studies,^19-21^ but currently there are insufficient genetic instruments to explore whether or not such associations are causal.

Coffee has been studied in a large number of observational studies of PD risk, and greater consumption has been shown to be associated with a reduced risk of PD.^7^ In this study, the point estimates from our main (IVW analysis) suggested a possible 12% increased relative risk of PD in those who drank coffee compared with those who did not, but the 95% confidence intervals were wide and included the null. If a true positive effect of 12% increase in PD risk exists, the present study would have been underpowered to detect this. Whilst this does not rule out a potential causal adverse role for coffee on PD risk, these results together with those of a recent randomised controlled trial of caffeine use in PD, which was terminated early because it failed to meet its primary outcome of motor symptom benefit,^22^ suggest that caffeine may neither prevent PD occurring nor be of benefit in those with the condition. However the early termination of this trial meant that the disease-modifying stage of the study never commenced and so empirical evidence for the role of coffee on PD progression is lacking. It is also important to note that potentially causal effects of coffee may not occur exclusively through caffeine, since coffee contains other chemicals (such as caffeic acid and polyphenols), which are also known to be biologically active.

If the overall effect of coffee consumption on PD risk is a true null, the link between coffee consumption and PD in observational studies requires explanation. One possibility is that being a “morning person” is causally associated with an increased risk of PD and that those that are not “morning persons” are more likely to consume coffee in greater quantities. In this scenario, a spurious protective association between coffee and PD could be induced through confounding by chronotype. Other factors, such as smoking, which is commonly associated with coffee drinking behaviour,^23^ could also confound the association.

### Circadian rhythm, caffeine, and Parkinson’s disease

Evidence from other sources also suggests that disruption of the circadian clock is associated with PD.^4,5^ Sleep disturbances such as REM sleep behaviour disorder and other non-motor features are common in PD and often arise several years before PD diagnosis.^24^ However, neuropathological studies have found that brain structures related to circadian rhythm are only affected in the later stage of PD, which suggests that disturbed circadian rhythm is not a consequence of neurodegeneration in PD.^25^ The evidence presented here suggests that the association may arise in the alternative direction, with altered circadian rhythm putting one at higher risk of PD. The mechanism by which this arises is not immediately apparent, but warrants further study. The circadian clock plays important roles in cellular homeostasis and under certain conditions may affect the way that neuronal cells prevent accumulation or clear aggregated proteins, which are the hallmark of neurodegenerative disease, including PD.^26^ An alternative explanation is the circadian rhythm is casually related to chronotype and PD via separate pathways, in which case intervention directed at chronotype would not necessary alter risk of PD.

As the most widely consumed psychostimulant agent globally, caffeine is known to promote wakefulness and induce or sustain performance.^27,28^ Experimental evidence (both *in vivo* and *in vitro*) suggests that caffeine delays the phase of the circadian clock measured at a cellular level and in human melatonin release cycle.^29^ Such alteration in circadian rhythm may explain why the point estimate for high coffee consumption was in the direction of increasing risk of PD, albeit the analysis was underpowered. Unfortunately, we were not able to further explore a causal relationship between coffee and chronotype using the available data.

### Strengths and limitations of this study

MR uses an IV approach to assess causal relationships between environmental exposures/intermediate phenotypes and disease outcomes. It relies on certain assumptions (summarised in the Methods section and supplementary material). Here we used two-sample MR to assess a causal effect of 1) liability towards being a “morning person” on risk of PD, 2) high coffee consumption on risk of PD, and 3) risk of PD on the probability of being a morning person. The methodological advantages of two-sample MR have been summarised previously^9^ and include: 1) cost and time saving, 2) access to very large sample sizes and resulting high statistical power, and 3) where bias does occur, it tends to be towards the null and estimates are conservative.^8^ Further advantages included the strength of the instruments (meaning that bias due to weak instruments was unlikely) and that we used a variety of MR analyses (each with different underlying assumptions) to confirm our findings. We also searched published literature both in humans and in fruit flies (*Drosophila*) to ascertain the relevance to the circadian clock for each of the SNPs we used to perform MR analyses (see supplementary material). In addition to the 11 loci deemed to play a plausible role in circadian rhythmicity by Hu and colleagues,^13^ this process suggested potential clock-control of an additional two genes in risk loci (13 out 15 total), indicating that the combined variants are an appropriate instrument for analyses relating to the circadian clock.

Finally, we took steps to ensure that our samples were not overlapping by using 23andMe GWAS data to identify the SNPs and the UK Biobank data to obtain the exposure estimates because controls from the PD GWAS may have also been used in the 23andMe discovery data. Separately, we also ensured that individuals used in a recent PD GWA study meta-analysis were not used in our reverse causation analysis. With the exception of the reverse causation analysis, our MR analyses used aggregated GWA study summary statistics from publicly available published data sets. Weaknesses of using aggregate data include the inability to consider and adjust for potential confounding factors.

In summary, we provide evidence to support a causal effect between liability towards being a “morning person” and PD. This implicates modulation of circadian rhythm as a potential therapeutic avenue to alter risk of PD.

**Table 1:**
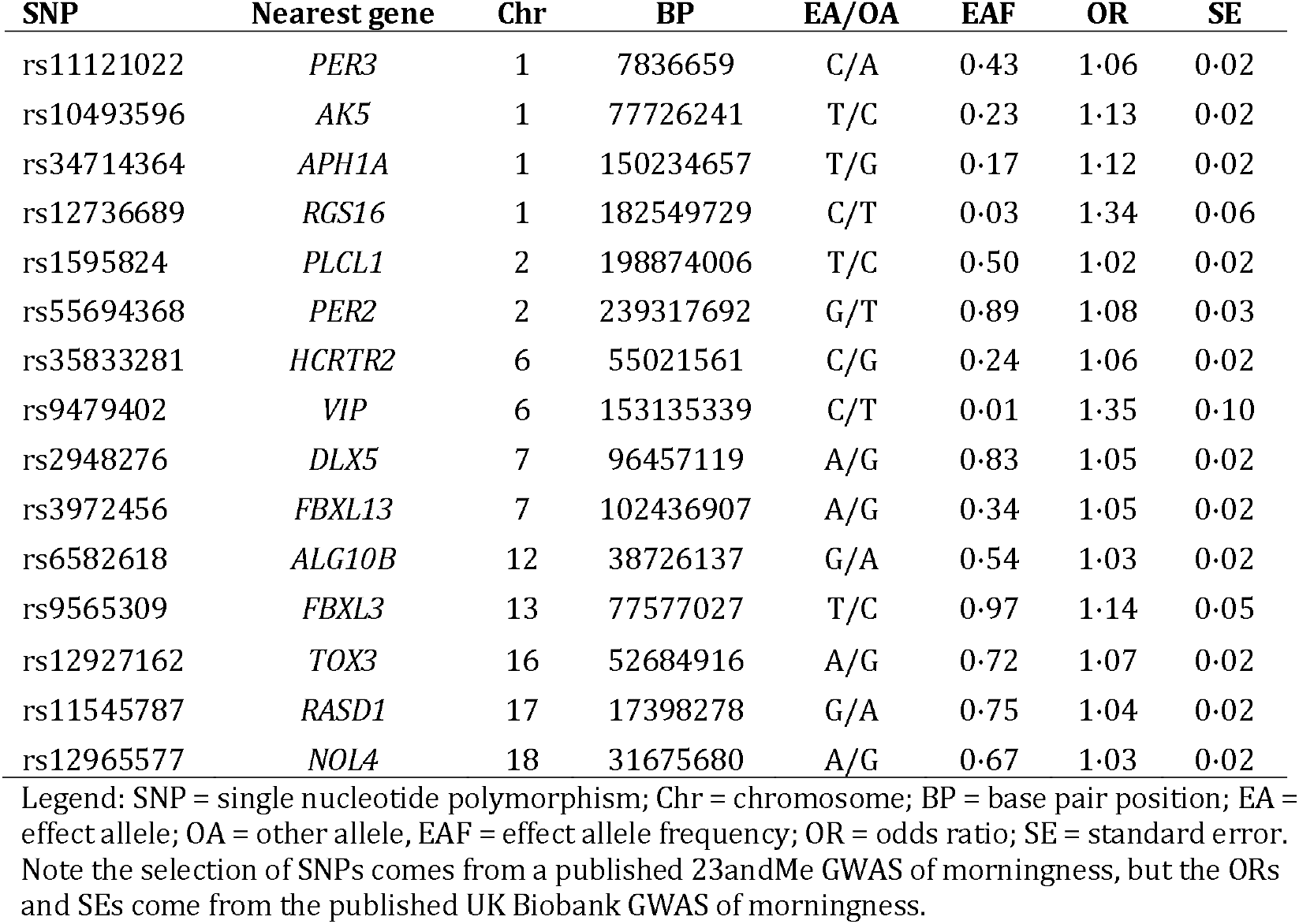
SNPs used to construct the morning person instrumental variable

**Table 2:**
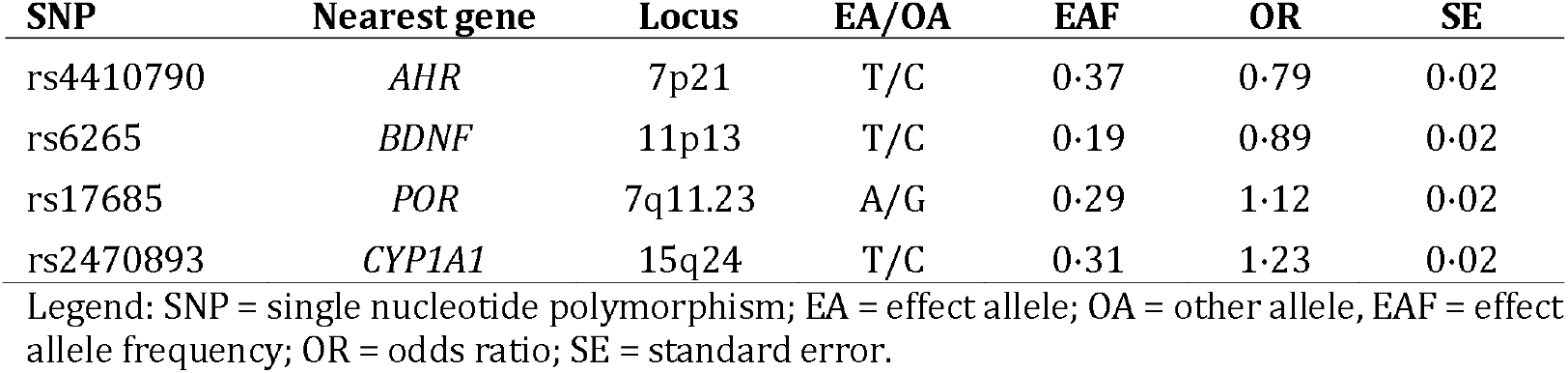
SNPs used to construct the coffee (high versus low/none) instrumental variable

## Declaration of interests

AJN is funded by a Barts and the London Charity Grant for Preventive Neurology. DAK is supported by an MB PhD Award from the International Journal of Experimental Pathology. JECJ is funded by an MRC New Investigator Award. GH, GDS and DAL work in a unit that receives funding from the University of Bristol and the UK Medical Research Council (MC_UU_1201/1 and MC_UU_1201/5). DAL is a National Institute of Health Research Senior Investigator (NF-SI-0166-10196). K Heilbron and DA Hinds are employees of 23andMe. MAN and AS participation in this study was supported in part by the Intramural Research Program of the National Institute on Aging, NIH. University College London Hospitals and University College London receive support from the Department of Health’s National Institute for Health Research (NIHR) Biomedical Research Centres (BRC). MAN is a paid consultant/contractor to the NIH. DAL reports grants from UK Medical Research Council, during the conduct of the study; grants from Medical Research council, grants from Wellcome Trust, grants from European Research Council, grants from Roche Diagnostics, grants from Medtronic, grants from Ferring Pharmaceuticals, outside the submitted work. NWW is an NIHR senior investigator and receives support from the JPND-MRC Comprehensive Unbiased Risk factor Assessment for Genetics and Environment in Parkinson’s disease (COURAGE). GH and JH report no relevant disclosures.

## Author Contributions

Conceptualisation & Design: AJN, DAK, JH, AS, MAN, NP, GDS, DAL, NWW. Acquisition and Analysis: AJN, DAK, KH, JECJ AS, DH, GDS, DAL, MAN. Interpretation: AJN, DAK, KH, JECJ, GH, DH, DAL, GDS, MAN, NWW. Writing – original draft: AJN, DAK, KH, JECJ, AS, MAN, NWW. Writing – review & editing: GH, JH, DH, DAL, GDS.

## Role of the funding source

The authors received no specific funding for this work. Therefore no funder had a role in study design, data collection and analysis, decision to publish, or preparation of the manuscript. The corresponding author should con rm that he or she had full access to all the data in the study and had nal responsibility for the decision to submit for publication

## Acknowledgements

Full acknowledgements and affiliations of IPDGC members are listed in the supplementary material. We would like to thank the research participants and employees of 23andMe for making this work possible. We also thank the following members of the 23andMe Research Team: Michelle Agee, Babak Alipanahi, Adam Auton, Robert K. Bell, Katarzyna Bryc, Sarah L. Elson, Pierre Fontanillas, Nicholas A. Furlotte, David A. Hinds, Karen E. Huber, Aaron Kleinman, Nadia K. Litterman, Jennifer C. McCreight, Matthew H. McIntyre, Joanna L. Mountain, Elizabeth S. Noblin, Carrie A.M. Northover, Steven J. Pitts, J. Fah Sathirapongsasuti, Olga V. Sazonova, Janie F. Shelton, Suyash Shringarpure, Chao Tian, Joyce Y. Tung, Vladimir Vacic, and Catherine H. Wilson.

## References

1. Lawlor DA, Harbord RM, Sterne JA, Timpson N, Davey Smith G. Mendelian randomization: using genes as instruments for making causal inferences in epidemiology. Stat Med 2008; 27(8): 1133–63.

2. Noyce AJ, Nalls MA. Mendelian Randomization – the Key to Understanding Aspects of Parkinson's Disease Causation? Mov Disord 2016; 31(4): 478–83.

3. Lane JM, Vlasac I, Anderson SG, et al. Genome-wide association analysis identifies novel loci for chronotype in 100,420 individuals from the UK Biobank. Nat Commun 2016; 7: 10889.

4. Breen DP, Vuono R, Nawarathna U, et al. Sleep and circadian rhythm regulation in early Parkinson disease. JAMA Neurol 2014; 71(5): 589–95.

5. De Pablo-Fernandez E, Breen DP, Bouloux PM, Barker RA, Foltynie T, Warner TT. Neuroendocrine abnormalities in Parkinson's disease. J Neurol Neurosurg Psychiatry 2017; 88(2): 176–85.

6. Chen H, Schernhammer E, Schwarzschild MA, Ascherio A. A prospective study of night shift work, sleep duration, and risk of Parkinson's disease. Am J Epidemiol 2006; 163(8): 726–30.

7. Noyce AJ, Bestwick JP, Silveira-Moriyama L, et al. Meta-analysis of early nonmotor features and risk factors for Parkinson disease. Ann Neurol 2012; 72(6): 893–901.

8. Pierce BL, Burgess S. Efficient design for Mendelian randomization studies: subsample and 2-sample instrumental variable estimators. Am J Epidemiol 2013; 178(7): 1177–84.

9. Lawlor DA. Commentary: Two-sample Mendelian randomization: opportunities and challenges. Int J Epidemiol 2016; 45(3): 908–15.

10. Bowden J, Davey Smith G, Burgess S. Mendelian randomization with invalid instruments: effect estimation and bias detection through Egger regression. Int J Epidemiol 2015; 44(2): 512–25.

11. Davey Smith G, Hemani G. Mendelian randomization: genetic anchors for causal inference in epidemiological studies. Hum Mol Genet 2014; 23(R1): R89–98.

12. Nalls MA, Pankratz N, Lill CM, et al. Large-scale meta-analysis of genome-wide association data identifies six new risk loci for Parkinson’s disease. Nat Genet 2014; 46(9): 989–93.

13. Hu Y, Shmygelska A, Tran D, Eriksson N, Tung JY, Hinds DA. GWAS of 89,283 individuals identifies genetic variants associated with self-reporting of being a morning person. Nat Commun 2016; 7: 10448.

14. Hemani G, Zheng J, Wade KH, et al. MR-Base: a platform for systematic causal inference across the phenome using billions of genetic associations. bioRxiv 2016.

15. Cornelis MC, Byrne EM, Esko T, et al. Genome-wide meta-analysis identifies six novel loci associated with habitual coffee consumption. Mol Psychiatry 2015; 20(5): 647–56.

16. Bowden J, Davey Smith G, Haycock PC, Burgess S. Consistent Estimation in Mendelian Randomization with Some Invalid Instruments Using a Weighted Median Estimator. Genet Epidemiol 2016; 40(4): 304–14.

17. Hartwig FP, Davey Smith G, Bowden J. Robust inference in summary data Mendelian randomization via the zero modal pleiotropy assumption International Journal of Epidemiology 2017.

18. Brion MJ, Shakhbazov K, Visscher PM. Calculating statistical power in Mendelian randomization studies. Int J Epidemiol 2013; 42(5): 1497–501. 19.

19. Todes CJ, Lees AJ. The pre-morbid personality of patients with Parkinson's disease. J Neurol Neurosurg Psychiatry 1985; 48(2): 97–100.

20. Heberlein I, Ludin HP, Scholz J, Vieregge P. Personality, depression, and premorbid lifestyle in twin pairs discordant for Parkinson's disease. J Neurol Neurosurg Psychiatry 1998; 64(2): 262–6.

21. Paulson GW, Dadmehr N. Is there a premorbid personality typical for Parkinson's disease? Neurology 1991; 41(5 Suppl 2): 73–6.

22. Postuma RB, Anang J, Pelletier A, et al. Caffeine as symptomatic treatment for Parkinson disease (Cafe-PD): A randomized trial. Neurology 2017.

23. Treur JL, Taylor AE, Ware JJ, et al. Associations between smoking and caffeine consumption in two European cohorts. Addiction 2016; 111(6): 1059–68.

24. Noyce AJ, Lees AJ, Schrag AE. The prediagnostic phase of Parkinson's disease. J Neurol Neurosurg Psychiatry 2016; 87(8): 871–8.

25. De Pablo-Fernandez E, Courtney R, Holton JL, Warner TT. Hypothalamic alpha-synuclein and its relation to weight loss and autonomic symptoms in Parkinson's disease. Mov Disord 2017; 32(2): 296–8.

26. Hastings MH, Goedert M. Circadian clocks and neurodegenerative diseases: time to aggregate? Curr Opin Neurobiol 2013; 23(5): 880–7.

27. Reyner LA, Horne JA. Early morning driver sleepiness: effectiveness of 200 mg caffeine. Psychophysiology 2000; 37(2): 251–6.

28. Bonnet MH, Gomez S, Wirth O, Arand DL. The use of caffeine versus prophylactic naps in sustained performance. Sleep 1995; 18(2): 97–104. 29.

29. Burke TM, Markwald RR, McHill AW, et al. Effects of caffeine on the human circadian clock in vivo and in vitro. Sci Transl Med 2015; 7(305): 305ra146.

