## Supplementary Materials for "Tendency towards being a “Morning person” increases risk of Parkinson’s disease: evidence from Mendelian randomisation"

**Noyce AJ et al**

Page 2 – Basic instrumental variable assumptions

Page 3 – Additional analyses

Page 5 – *Drosophila* homologue pipeline

Page 6 – Table 1: Genes found at risk loci for chronotype and relevance to the circadian clock

Page 8 – Table 2: Cohorts involved included in the IPDGC meta GWAS

Page 10 – IPDGC members and affiliations

Page 12 – IPDGC acknowledgements

**Basic instrumental variable (IV) assumptions**

Mendelian randomisation (MR) relies on certain assumptions, which are listed here:

1. **The IV must be robustly associated with the exposure of interest.** Calculating an F statistic and R^2^ value can be used to check this assumption. The IV may only explain tiny amount of variance in the exposure (R^2^) and therefore studies often have to be large. As a result of genome-wide association (GWA) studies, there are increasing numbers of genetic variants that account for the variance in a range of exposures and outcomes, allowing instruments of greater strength to be constructed.
2. **The IV must be independent of known confounders.** In MR studies that have individual level data, one can check for known confounders and compare the frequency of these between the two levels of the IV. In two-sample MR, the absence of individual level data about potential confounders can hamper the ability to test this assumption.
3. **The IV must be independent of the outcome, given the exposure and confounders.** In other words, there must be no alternative path, other than via the exposure, that the IV influences the outcome. This is known as the exclusion restriction criteria. One situation that violates this assumption is horizontal pleiotropy, in which there are alternative pathways through which the IV affects the outcome.

**Additional analyses**

We undertook a variety of additional analyses to confirm the veracity of our main observations:

1. Two sample MR of liability to being a “morning person” (using variants, effect estimates and standard errors from Hu et al, 2016) and risk of Parkinson’s disease (using IPDGC data).

The main analysis was repeated using data only from 23andMe (Hu et al, 2016). The results from IVW analysis were similar to the results from the main analysis, which used effect estimates and standard errors from UK Biobank, (Lane et al, 2016) and suggested that liability towards being a “morning person” was causally related to risk of PD (OR 1.16; 95% CI 1.03-1.30). There was no evidence of heterogeneity (Q=13.2, p=0.513, I^2^=0%). The MR-Egger analysis did not suggest the IVW estimate was biased by net directional horizontal pleiotropy OR 1.23 (95% CI 0.93-1.53), intercept -0.006 (p=0.67).

2. Two sample MR of liability to being a “morning person” (using variants listed by Hu et al (2016), and effect estimates and standard errors for these variants from the full unpublished summary statistics of the chronotype GWAS from UK Biobank (Broad Institute website)) and risk of Parkinson’s disease (using IPDGC data).

The main analysis was repeated using unpublished summary statistics (effect size estimates and standard errors) from the full UK Biobank GWAS of chronotype, again for the variants identified from 23andMe (Hu et al, 2016). Summary statistics were available for 14 of the 15 SNPs reported by Hu et al (2016). rs3972456 was not available and there was no identifiable proxy. Therefore 14 SNPs were used in the combined instrument.

The results from IVW analysis showed that liability towards being a “morning person” was causally related to risk of PD (OR 1.80; 95% CI 1.05-3.90; p=0.034). There was no evidence of heterogeneity (Q=12.5, p=0.485, I^2^=0%). The MR-Egger analysis did not suggest the IVW estimate was biased by net directional horizontal pleiotropy OR 2.29 (95% CI 0.67-7.80; p=0.166), intercept -0.006 (p=0.642). These results are presented in the figure below.

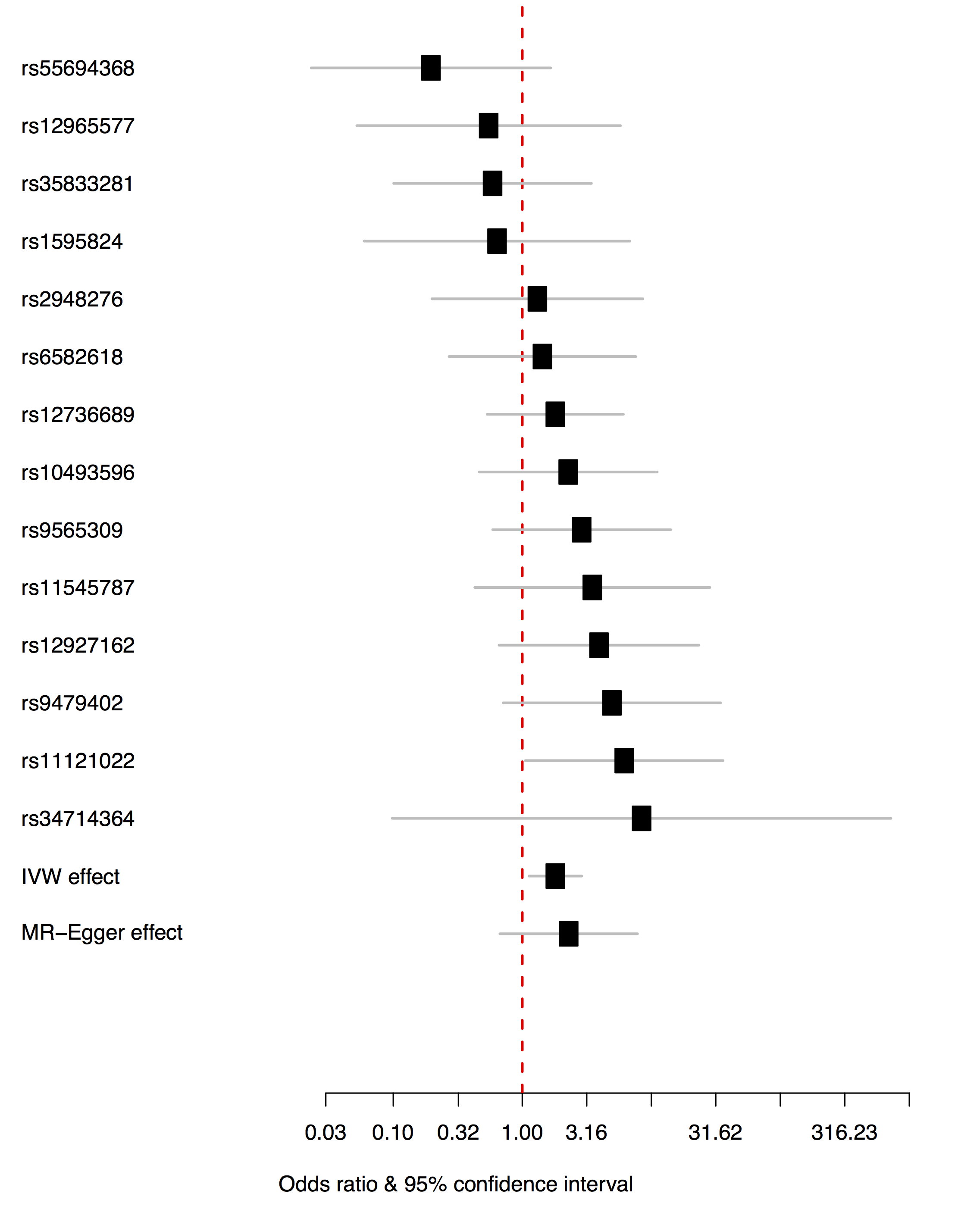

Legend: Forest plot shows the per-variant odds ratio and 95% confidence interval for liability towards being a “morning person” and risk of PD. Combined estimates are given for inverse weight variance and MR-Egger.

3. Two sample MR of liability towards being a “morning person” using a novel instrument derived from the full unpublished summary statistics of the chronotype GWAS from UK Biobank (Broad Institute website) and risk of Parkinson’s disease using a) IPDGC data and b) 23andMe data.
   a. 71 SNPs were independently associated with chronotype over a four point ordinal scale from extreme “evening person” to extreme “morning person” (i.e. different to the binary case control analysis undertaken in published UK Biobank and 23andMe data in both the direction of measurement and the number of levels of the exposure). This instrument gave rise to a null result for the association between chronotype and PD risk using IPDGC data (IVW OR 1.1, SE 0.123, p-value=0.426).
   b. The same instrument yielded a null result in 23andMe PD case/control data (p-value=0.877).
   c. We explored the reasons why results may be attenuated using the new instrument generated from the full unpublished UK Biobank chronotype GWAS, which include:

- ‘Winner’s curse’ which will bias the effect estimate in two sample MR towards the null.
- Two sample MR in general tends to bias effect estimates towards the null.
- There are many more SNPs at marginal significance in the unpublished UK Biobank Chronotype GWAS and also the possibility of greater net horizontal pleiotropic effect.
- The betas in the unpublished full UK Biobank chronotype GWAS were derived from regression on an ordinal exposure, where original instruments previously compared extremes, therefore betas are smaller, and furthermore the exposure-outcome relationship may not be linear.

4. One sample MR of liability towards being a “morning person” and risk of Parkinson’s disease (in 23andMe data exclusively) using a genetic risk score.

Using 23andMe data only, a new GWAS of liability towards being a “morning person” identified over 200 SNPs and the MR effect was in favour of causal association between tendency towards being a morning person and risk of PD (p=0.03). In 23andMe the observational study association between chronotype and risk of PD was a null and the effect of this in one sample MR would be expected to bias the result towards null (i.e. this is an under-estimate). These findings support the main analyses using the previously published 15 variant chronotype instrument (published by Hu et al, 2016).

5. LD regression between the two GWASs (UK Biobank chronotype GWAS and the 23andMe chronotype) found that they were highly correlated (r=0.93). This suggests that they are measuring the same trait, but on a different scale, as discussed above.

***Drosophila* homologue pipeline**

Given that four of the 15 variants identified from the 23andMe chronotype GWAS (Hu et al, 2016) had no known link to the circadian clock, either in humans or small animal models, we undertook a further in silico search of literature pertaining to the *Drosophila* circadian clock, which has been extensively studied at the molecular level. We assessed whether any of the homologs of the loci identified by Hu et al. (2016) were either components of, or were regulated by, the circadian clock. This yielded evidence for genes in two more of the reported loci, bringing the total number of genes with a plausible link to the circadian clock to 13 out of 15 (see table below). The pipeline used for identification of *Drosophila* homologs and potential functions was as follows.

1. The closest *Drosophila* homolog of each loci based on amino-acid sequence was identified using BLAST. A reverse BLAST (i.e using the *Drosophila* protein to screen the human proteome) was also used to confirm orthology, paralogy or a more distant homology between the proteins of interest.
2. Once a *Drosophila* homologue (if present) was identified, five different sources were used to identify links to the circadian clock:
   a. Research papers documented on Flybase (http://flybase.org)
   b. Pubmed, using the search terms 'gene x' + *Drosophila* + either 'circadian' or 'clock'.
   c. Abruzzi et al., (2011) *Genes and Development*: a list of *Drosophila* genes with cycling binding of the CLOCK transcription factor.
   d. Nagoshi et al., (2009) *Nature Neuroscience*: a list of genes whose transcription is enriched in clock neurons in the *Drosophila* brain.
   e. Abruzzi et al., (2017) *PLoS Genetics*: a list of *Drosophila* genes whose transcription cycles in one of three groups of clock neurons (LNv, LNd and DN1) in the *Drosophila* brain.

**Table 1: Genes in risk loci for chronotype (adapted from Hu et al, 2016) and corresponding *Drosophila* homologues. Functions or patterns of regulation potentially linked to the circadian clock are described, if known.**

|  | **Human gene/SNP/ Chromosome** | **Potential circadian role and relevant model (as reported by Hu et al, 2016)** | ***Drosophila* homologue** | **Molecular function related to the circadian clock?** | **Link to circadian clock in literature?** | **Evidence of a direct role in clock function?** | **Evidence of regulation by the clock?** |
| --- | --- | --- | --- | --- | --- | --- | --- |
| **Genes with known circadian roles** | RGS16  rs12736689  Chr 1 | RGS16 knock-out mice have longer circadian period | *double hit* (*dhit*) | N/A | N/A | N/A | N/A |
|  | VIP  rs9479402  Chr 6 | VIP encodes neuropeptide which, when injected, prolongs REM sleep in rabbits | N/A | N/A | N/A | N/A | N/A |
|  | PER2  rs55694368  Chr 2 | Associated with human familial advanced sleep phase syndrome | *period* | Circadian transcription factor | Yes | Yes | N/A |
|  | HCRTR2  rs35833281  Chr 6 | Mutations in HCRTR2 linked to narcolepsy in dogs and humans | N/A | N/A | N/A | N/A | N/A |
|  | RASD1  rs11545787  Chr 17 | Deletion of RASD1 leads to reduction of photic entrainment in mice | cg8641 | N/A | N/A | N/A | N/A |
|  | PER3  rs1121022  Chr 1 | Affects the sensitivity of the circadian system to light26 and is involved in sleep/wake activity. | *period* | Circadian transcription factor | Yes | Yes | N/A |
|  | FBXsdsdL3  rs9565309  Chr 13 | Mutant FBXL3 mice were shown to have an extended circadian period. | *partner of paired* (*ppa*) | N/A | N/A | N/A | N/A |
| **Genes with plausible circadian role** | PLCL1  rs1595824  Chr 2 | Expressed in CNS and binds to the GABA  type A receptor | *norpA* | Provides light input to circadian clock neurons via photoreceptors | Yes | Yes | N/A |
|  | APH1A  rs34714364  Chr 1 | Part of the gamma-secretase complex, which cleaves beta-APP & regulated by orexin and sleep-wake cycle | *aph-1* | N/A | N/A | N/A | N/A |
|  | FBXL13  rs3972456  Chr 7 | Protein-ubiquitin ligase and may have a circadian role similar to FBXL3 | cg9003 | N/A | N/A | N/A | N/A |
|  | NOL4  rs12965577  Chr 18 | Significant change in expression in mice with knock in alpha-1 subunit of GABA(A) receptor | N/A | N/A | N/A | N/A | N/A |
| **Genes with less clear circadian role** | TOX3  rs12927162  Chr 16 | Associated with restless leg syndrome | N/A | N/A | N/A | N/A | N/A |
|  | AK5  rs10493596  Chr 1 | Regulates adenine nucleotide metabolism in the brain | *adk1* | N/A | Yes | No | Target of CLOCK binding in fly brain and shows evidence of cycling of both CLOCK and POLII binding (Abruzzi et al., (2011) Genes Dev |
|  | DLX5  rs2948276  Chr 7 | Associated with hand and foot malformations | *distal-less* (*dll*) | N/A | Yes | No | Transcript cycles in DN1p clock cells (Abruzzi et al. (2017) PLoS Genetics |
|  | ALG10B  rs6582618  Chr 12 | Has a role in cardiac rhythm | *alg10* | N/A | N/A | N/A | N/A |

**References for *Drosophila* homologue pipeline**

1. Abruzzi KC, Zadina A, Luo W, et al. RNA-seq analysis of *Drosophila* clock and non-clock neurons reveals neuron-specific cycling and novel candidate neuropeptides. PLoS Genetics 2017 3(2): e1006613
2. Abruzzi KC, Rodriquez J, Menet JS, et al. *Drosophila* CLOCK target gene characterization: implications for circadian tissue-specific gene expression. Genes & Dev. 2011. 25: 2374-2386
3. Nagoshi E, Sugino K, Kula E, et al. Dissecting differential gene expression within the circadian neuronal circuit of *Drosophila.* Nature Neuroscience 2010;13:60–68

**Table 2: Cohorts included in the IPDGC meta-analysis (adapted from Nalls et al. Nat Genet 2014; 46 : 989–993).**

| **Study (Country)** | **Cases (N)** | **Controls (N)** | **Total sample size (N)** | **Cases % female** | **Controls % female** | **Cases, age at onset, mean (SD)** | **Controls, age at ascertainment, mean (SD)** | **Case ascertainment criteria** | **Control ascertainment criteria** |
| --- | --- | --- | --- | --- | --- | --- | --- | --- | --- |
| IPDGC (Iceland, deCODE) | 604 | 4916 | 5520 | 47.8 | 56.9 | 62.2 (12.3) | 80.2 (7.5) | Clinic visit; standard UK Brain Bank criteria with a modification to allow the inclusion of cases that had a family history of PD | Absence of self report, ICD-9, and medications |
| IPDGC (France) | 985 | 1984 | 2969 | 41.2 | 33 | 48.9 (12.8) | 73.7 (5.4) | Clinic visit; standard UK Brain Bank criteria with a modification to allow the inclusion of cases that had a family history of PD | Clinic visit and self-report |
| IPDGC (Germany) | 667 | 937 | 1604 | 39.8 | 48 | 55.7 (11.5) | 47.4 (12.4) | Clinic visit; standard UK Brain Bank criteria with a modification to allow the inclusion of cases that had a family history of PD | Population controls |
| IPDGC (Netherlands) | 744 | 2019 | 2763 | 36 | 63.9 | 55.6 (11.8) | 55.7 (5.8) | Clinic visit; standard UK Brain Bank criteria with a modification to allow the inclusion of cases that had a family history of PD | Population controls |
| IPDGC (USA) | 937 | 1896 | 2833 | 40.5 | 52.8 | 57.8 (13.2) | 63.3 (10.1) | Clinic visit; standard UK Brain Bank criteria with a modification to allow the inclusion of cases that had a family history of PD | Clinic visit and self-report |
| IPDGC (UK) | 1705 | 5200 | 6905 | 43.3 | 49.5 | 64.2 (12.4) | 53 (0) | Clinic visit; standard UK Brain Bank criteria with a modification to allow the inclusion of cases that had a family history of PD | Population controls |
| 23andMe.v2 (USA, Europe) | 3261 | 29499 | 32760 | 39.5 | 41.6 | 64.2 (11.2) | 49.1 (14.9) | Self-report of PD diagnosis by clinician | Self-report of neurological disease, memory loss, tremor or family history of PD removed |
| 23andMe.v3 (USA, Europe) | 866 | 32538 | 33424 | 39 | 39.6 | 63.9 (10.9) | 43.5 (15.9) | Self-report of PD diagnosis by clinician | Self-report of neurological disease, memory loss, tremor or family history of PD removed |
| Ashkenazi Jewish (USA) | 268 | 178 | 446 | 33.2 | 57.3 | 59.9 (12.1) | 69.8 (8.8) | Clinic visit; standard UK Brain Bank criteria with a modification to allow the inclusion of cases that had a family history of PD | Medical records and self-report |
| HIHG (USA) | 574 | 619 | 1193 | 36.9 | 65.4 | 57.2 (12.03) | 69.3 (9.9) | Clinic visit; standard UK Brain Bank criteria with a modification to allow the inclusion of cases that had a family history of PD | Clinic visit and self-report |
| NGRC (USA) | 1956 | 1982 | 3938 | 33.2 | 61.3 | 58.6 (11.7) | 70.3 (14.1) | Clinic visit; standard UK Brain Bank criteria with a modification to allow the inclusion of cases that had a family history of PD | Clinic visit and self-report |
| PROGENI-GenePD (USA) | 828 | 852 | 1680 | 40.1 | 60.2 | 62.1 (10.7) | 54.9 (13.1) | Clinic visit; standard UK Brain Bank criteria with a modification to allow the inclusion of cases that had a family history of PD | Clinic visit and self-report |
| CHARGE-CHS (USA) | 107 | 3164 | 3271 | 44.9 | 61.4 | 73 (5.1) | 72.3 (5.4) | Self-report, ICD-9, and medications | Absence of self report, ICD-9, and medications |
| CHARGE-FHS (USA) | 60 | 3889 | 3949 | 41.7 | 54.6 | 76.2 (10.8) | 64.2 (12.2) | Clinic visit; standard UK Brain Bank criteria with a modification to allow the inclusion of cases that had a family history of PD | On-going surveillance and periodic physician examination |
| CHARGE-AGES-RS (Iceland) | 146 | 5609 | 5755 | 54.8 | 59 | 75 (8.2) | 69 (8.9) | Clinic visit; medication use; ICD codes | Absence of self report, ICD-9, and medications |

**IPDGC consortium members and affiliations:**

**United Kingdom:** Alastair J Noyce (Preventive Neurology Unit, Wolfson Institute of Preventive Medicine, QMUL, London, UK and Department of Molecular Neuroscience, UCL, London, UK), Arianna Tucci (Department of Molecular Neuroscience, UCL Institute of Neurology, London, UK), Demis A Kia (UCL Genetics Institute; and Department of Molecular Neuroscience, UCL Institute of Neurology, London, UK), Gavin Charlesworth (Department of Molecular Neuroscience, UCL Institute of Neurology, London, UK), Manuela Tan (Department of Clinical Neuroscience, University College London, London, UK), Henry Houlden (Department of Molecular Neuroscience, UCL Institute of Neurology, London, UK),  Huw R Morris (Department of Clinical Neuroscience, University College London, London, UK), Helene Plun-Favreau (Department of Molecular Neuroscience, UCL Institute of Neurology, London, UK), Peter Holmans (Biostatistics & Bioinformatics Unit, Institute of Psychological Medicine and Clinical Neuroscience, MRC Centre for Neuropsychiatric Genetics & Genomics, Cardiff, UK), John Hardy (Department of Molecular Neuroscience, UCL Institute of Neurology, London, UK), Jose M Bras (Department of Molecular Neuroscience, UCL Institute of Neurology, London, UK), John Quinn (Institute of Translational Medicine, University of Liverpool, Liverpool, UK), Kin Y Mok (Department of Molecular Neuroscience, UCL Institute of Neurology, London, UK), Kimberley Billingsley (Institute of Translational Medicine, University of Liverpool, Liverpool, UK), Nicholas W Wood (UCL Genetics Institute; and Department of Molecular Neuroscience, UCL Institute of Neurology, London, UK), Patrick Lewis (University of Reading, Reading, UK), , Rita Guerreiro (Department of Molecular Neuroscience, UCL Institute of Neurology, London, UK), Ruth Lovering (University College London, London, UK), Raquel Duran Ogalla (University College London, London, UK), Lea R’Bibo (Department of Molecular Neuroscience, UCL Institute of Neurology, London, UK), Mina Ryten (Department of Molecular Neuroscience, UCL Institute of Neurology, London, UK), Valentina Escott-Price (MRC Centre for Neuropsychiatric Genetics and Genomics, Cardiff University School of Medicine, Cardiff, UK), Viorica Chelban (Department of Molecular Neuroscience, UCL Institute of Neurology, London, UK), Thomas Foltynie (UCL Institute of Neurology, London, UK), Una-Marie Sheerin (Department of Molecular Neuroscience, UCL Institute of Neurology, London, UK), Nigel Williams (MRC Centre for Neuropsychiatric Genetics and Genomics, Cardiff, UK),

**France:** Alexis Brice (Institut du Cerve au et de la Moelle épinière, ICM, Inserm U 1127, CNRS, UMR 7225, Sorbonne Universités, UPMC University Paris 06, UMR S 1127, AP-HP, Pitié-Salpêtrière Hospital, Paris, France), Fabrice Danjou (Institut du Cerveau et de la Moelle épinière, ICM, Inserm U 1127, CNRS, UMR 7225, Sorbonne Universités, UPMC University Paris 06, UMR S 1127, AP-HP, Pitié-Salpêtrière Hospital, Paris, France), Suzanne Lesage (Institut du Cerveau et de la Moelle épinière, ICM, Inserm U 1127, CNRS, UMR 7225, Sorbonne Universités, UPMC University Paris 06, UMR S 1127, AP-HP, Pitié-Salpêtrière Hospital, Paris, France), Jean-Christophe Corvol (Institut du Cerveau et de la Moelle épinière, ICM, Inserm U 1127, CNRS, UMR 7225, Sorbonne Universités, UPMC University Paris 06, UMR S 1127, Centre d’Investigation Clinique Pitié Neurosciences CIC-1422, AP-HP, Pitié-Salpêtrière Hospital, Paris, France), Maria Martinez (INSERM UMR 1220; and Paul Sabatier University, Toulouse, France),

**Germany:** Anamika Giri (Department for Neurodegenerative Diseases, Hertie Institute for Clinical Brain Research, University of Tübingen, and DZNE, German Center for Neurodegenerative Diseases, Tübingen, Germany), Angelika Oehmig (Department for Neurodegenerative Diseases, Hertie Institute for Clinical Brain Research, University of Tübingen, and DZNE, German Center for Neurodegenerative Diseases, Tübingen, Germany), Claudia Schulte (Department for Neurodegenerative Diseases, Hertie Institute for Clinical Brain Research), Kathrin Brockmann (Department for Neurodegenerative Diseases, Hertie Institute for Clinical Brain Research, University of Tübingen, and DZNE, German Center for Neurodegenerative Diseases, Tübingen, Germany), Javier Simón-Sánchez (Department for Neurodegenerative Diseases, Hertie Institute for Clinical Brain Research, University of Tübingen, and DZNE, German Center for Neurodegenerative Diseases, Tübingen, Germany), Peter Heutink (DZNE, German Center for Neurodegenerative Diseases and Department for Neurodegenerative Diseases, Hertie Institute for Clinical Brain Research, University of Tübingen, Tübingen, Germany), Patrizia Rizzu  (DZNE, German Center for Neurodegenerative Diseases), Manu Sharma (Centre for Genetic Epidemiology, Institute for Clinical Epidemiology and Applied Biometry, University of Tubingen and Department for Neurodegenerative Diseases, Hertie Institute for Clinical Brain Research, University of Tübingen Germany), Thomas Gasser (Department for Neurodegenerative Diseases, Hertie Institute for Clinical Brain Research, and DZNE, German Center for Neurodegenerative Diseases, Tübingen, Germany),

**United States of America:** Aude Nicolas (Laboratory of Neurogenetics, National Institute on Aging, Bethesda, MD, USA), Mark R Cookson (Laboratory of Neurogenetics, National Institute on Aging, Bethesda, USA), Sara Bandres-Ciga  (Laboratory of Neurogenetics, National Institute on Aging, Bethesda, MD, USA), Cornelis Blauwendraat (National Institute on Aging and National Institute of Neurological Disorders and Stroke, USA), Faraz Faghri (Laboratory of Neurogenetics, National Institute on Aging, Bethesda, USA; Department of Computer Science, University of Illinois at Urbana-Champaign, Urbana, IL, USA), J Raphael Gibbs (Laboratory of Neurogenetics, National Institute on Aging, Bethesda, MD, USA), Dena G Hernandez (Laboratory of Neurogenetics, National Institute on Aging, Bethesda, MD, USA), Joshua M. Shulman (Baylor College of Medicine, Houston, Texas, USA), Mike A. Nalls (Laboratory of Neurogenetics, National Institute on Aging, Bethesda, USA; CEO/Consultant Data Tecnica International, Glen Echo, MD, USA), Laurie Robak (Baylor College of Medicine, Houston, Texas, USA), Steven Lubbe (Ken and Ruth Davee Department of Neurology, Northwestern University Feinberg School of Medicine, Chicago, IL, USA), Steven Finkbeiner (Departments of Neurology and Physiology, University of California, San Francisco; Gladstone Institute of Neurological Disease; Taube/Koret Center for Neurodegenerative Disease Research, San Francisco, CA, USA),  Niccolo E. Mencacci (Northwestern University Feinberg School of Medicine, Chicago, IL, USA), Codrin Lungu (National Institutes of Health Division of Clinical Research, NINDS, National Institutes of Health, Bethesda, MD, USA), Andrew B Singleton (Laboratory of Neurogenetics, National Institute on Aging, Bethesda, MD, USA), Sonja Scholz (Neurodegenerative Diseases Research Unit, National Institute of Neurological Disorders and Stroke, Bethesda, MD, USA), Xylena Reed (Laboratory of Neurogenetics, National Institute on Aging, Bethesda, MD, USA).

**Canada:** Ziv Gan-Or (Montreal Neurological Institute and Hospital, Department of Neurology & Neurosurgery, Department of Human Genetics, McGill University, Montréal, QC, H3A 0G4, Canada), Guy A. Rouleau (Montreal Neurological Institute and Hospital, Department of Neurology & Neurosurgery, Department of Human Genetics, McGill University, Montréal, QC, H3A 0G4, Canada)

**The Netherlands:** Jacobus J van Hilten (Department of Neurology, Leiden University Medical Center, Leiden, Netherlands), Johan Marinus (Department of Neurology, Leiden University Medical Center, Leiden, Netherlands)

**Spain:** Juan A. Botía (Universidad de Murcia, Murcia, Spain), Jordi Clarimón (Memory Unit, Department of Neurology, IIB Sant Pau, Hospital de la Santa Creu i Sant Pau, Universitat Autònoma de Barcelona, Barcelona, and Centro de Investigación Biomédica en Red en Enfermedades Neurodegenerativas (CIBERNED), Madrid), Oriol Dols-Icardo (Memory Unit, Department of Neurology, IIB Sant Pau, Hospital de la Santa Creu i Sant Pau, Universitat Autònoma de Barcelona, Barcelona, and Centro de Investigación Biomédica en Red en Enfermedades Neurodegenerativas (CIBERNED), Madrid), Jaime Kulisevsky (Movement Disorders Unit, Department of Neurology, IIB Sant Pau, Hospital de la Santa Creu i Sant Pau, Universitat Autònoma de Barcelona, Barcelona, and Centro de Investigación Biomédica en Red en Enfermedades Neurodegenerativas (CIBERNED)), Javier Pagonabarraga (Movement Disorders Unit, Department of Neurology, IIB Sant Pau, Hospital de la Santa Creu i Sant Pau, Universitat Autònoma de Barcelona, Barcelona, and Centro de Investigación Biomédica en Red en Enfermedades Neurodegenerativas (CIBERNED)), Juan Marín (Movement Disorders Unit, Department of Neurology, IIB Sant Pau, Hospital de la Santa Creu i Sant Pau, Universitat Autònoma de Barcelona, Barcelona, and Centro de Investigación Biomédica en Red en Enfermedades Neurodegenerativas (CIBERNED)).

**Norway:** Lasse Pihlstrom (Department of Neurology, Oslo University Hospital, Oslo, Norway)

Estonia: Sulev Koks (Department of Pathophysiology, University of Tartu, Tartu, Estonia), Pille Taba (Department of Neurology and Neurosurgery, University of Tartu, Tartu, Estonia)

**IPDGC ACKNOWLEDGMENTS:**

We would like to thank all of the subjects who donated their time and biological samples to be a part of this study. This work was supported in part by the Intramural Research Programs of the National Institute of Neurological Disorders and Stroke (NINDS), the National Institute on Aging (NIA), and the National Institute of Environmental Health Sciences both part of the National Institutes of Health, Department of Health and Human Services; project numbers 1ZIA-NS003154, Z01-AG000949-02 and Z01-ES101986. In addition this work was supported by the Department of Defense (award W81XWH-09-2-0128), and The Michael J Fox Foundation for Parkinson’s Research. This work was supported by National Institutes of Health grants R01NS037167, R01CA141668, P50NS071674, American Parkinson Disease Association (APDA); Barnes Jewish Hospital Foundation; Greater St Louis Chapter of the APDA; Hersenstichting Nederland; the Prinses Beatrix Fonds. The KORA (Cooperative Research in the Region of Augsburg) research platform was started and financed by the Forschungszentrum für Umwelt und Gesundheit, which is funded by the German Federal Ministry of Education, Science, Research, and Technology and by the State of Bavaria. This study was also funded by the National Genome Research Network (NGFNplus number 01GS08134, German Federal Ministry for Education and Research), the German Federal Ministry of Education and Research (NGFN 01GR0468, PopGen) and 01EW0908 in the frame of ERA-NET NEURON. In addition, this work was supported by the EU Joint Programme - Neurodegenerative Diseases Research (JPND) project under the aegis of JPND ([www.jpnd.eu](http://www.jpnd.eu)) through Germany, BMBF, funding code 01ED1406 and iMed - the Helmholtz Initiative on Personalized Medicine. The French GWAS work was supported by the French National Agency of Research (ANR-08-MNP-012).  This study was also funded by France-Parkinson Association, the French program “Investissements d’avenir” funding (ANR-10-IAIHU-06) and a grant from Assistance Publique-Hôpitaux de Paris (PHRC, AOR-08010) for the French clinical data.  This study was also sponsored by the Landspitali University Hospital Research Fund (grant to SSv); Icelandic Research Council (grant to SSv); and European Community Framework Programme 7, People Programme, and IAPP on novel genetic and phenotypic markers of Parkinson’s disease and Essential Tremor (MarkMD), contract number PIAP-GA-2008-230596 MarkMD (to HP and JHu).  The McGill study was funded by the Michael J. Fox Foundation and the Canadian Consortium on Neurodegeneration in Aging (CCNA). This study utilized the high-performance computational capabilities of the Biowulf Linux cluster at the National Institutes of Health, Bethesda, Md. (http://biowulf.nih.gov), and DNA panels, samples, and clinical data from the National Institute of Neurological Disorders and Stroke Human Genetics Resource Center DNA and Cell Line Repository. People who contributed samples are acknowledged in descriptions of every panel on the repository website. We thank the French Parkinson’s Disease Genetics Study Group and the Drug Interaction with genes (DIGPD) study group: Y Agid, M Anheim, A-M Bonnet, M Borg, A Brice, E Broussolle, J-C Corvol, P Damier, A Destée, A Dürr, F Durif, A Elbaz, D Grabil, S Klebe, P. Krack, E Lohmann, L. Lacomblez, M Martinez, V Mesnage, P Pollak, O Rascol, F Tison, C Tranchant, M Vérin, F Viallet, and M Vidailhet. We also thank the members of the French 3C Consortium: A  Alpérovitch, C Berr, C Tzourio, and P Amouyel for allowing us to use part of the 3C cohort, and D Zelenika for support in generating the genome-wide molecular data. We thank P Tienari (Molecular Neurology Programme, Biomedicum, University of Helsinki),  T Peuralinna (Department of Neurology, Helsinki University Central Hospital), L Myllykangas (Folkhalsan Institute of Genetics and Department of Pathology, University of Helsinki), and R Sulkava (Department of Public Health and General Practice Division of Geriatrics, University of Eastern Finland) for the Finnish controls (Vantaa85+ GWAS data).  We used genome-wide association data generated by the Wellcome Trust Case-Control Consortium 2 (WTCCC2) from UK patients with Parkinson’s disease and UK control individuals from the 1958 Birth Cohort and National Blood Service. Genotyping of UK replication cases on ImmunoChip was part of the WTCCC2 project, which was funded by the Wellcome Trust (083948/Z/07/Z). UK population control data was made available through WTCCC1. This study was supported by the Medical Research Council and Wellcome Trust disease centre (grant WT089698/Z/09/Z to NW, JHa, and ASc). As with previous IPDGC efforts, this study makes use of data generated by the Wellcome Trust Case-Control Consortium. A full list of the investigators who contributed to the generation of the data is available from www.wtccc.org.uk. Funding for the project was provided by the Wellcome Trust under award 076113, 085475 and 090355.  This study was also supported by Parkinson’s UK (grants 8047 and J-0804) and the Medical Research Council (G0700943 and G1100643). We thank Jeffrey Barrett and Jason Downing for assistance with the design of the ImmunoChip and NeuroX arrays. DNA extraction work that was done in the UK was undertaken at University College London Hospitals, University College London, who received a proportion of funding from the Department of Health’s National Institute for Health Research Biomedical Research Centres funding. This study was supported in part by the Wellcome Trust/Medical Research Council Joint Call in Neurodegeneration award (WT089698) to the Parkinson’s Disease Consortium (UKPDC), whose members are from the UCL Institute of Neurology, University of Sheffield, and the Medical Research Council Protein Phosphorylation Unit at the University of Dundee. We thank the Quebec Parkinson’s Network (http://rpq-qpn.org) and its members. Mike A. Nalls’ participation is supported by a consulting contract between Data Tecnica International and the National Institute on Aging, NIH, Bethesda, MD, USA, as a possible conflict of interest Dr. Nalls also consults for Illumina Inc, the Michael J. Fox Foundation, University of California Healthcare and Genoom Health among others.  This work was supported by the Medical Research Council grant MR/N026004/1.
